# Liproxstatin-1 alleviates cartilage degradation by inhibiting chondrocyte ferroptosis in temporomandibular joint

**DOI:** 10.1101/2023.04.10.536321

**Authors:** Bei Cheng, Jun Zhang, Qinhao Shen, Zheyi Sun, Yingwei Luo, Yu Hu

## Abstract

Ferroptosis contribute to temporomandibular joint osteoarthritis (TMJOA) lesion development is still poorly understood. In this study, we used different TMJOA animal models to detect whether ferroptosis is related to onset of TMJOA which modelling by monosodium iodoacetate (MIA), IL-1β, occlusion disorder (OD) and unilateral anterior crossbite (UAC). Immunohistochemical staining and Western blot analysis were used to detect ferroptosis proteins and cartilage degradation related protein expression. Our results revealed that lower level of ferroptosis-related proteins GPX4 in cartilage layer, but the level of ACSL4 and P53 increase in that of condyle. Injection of ferroptosis inhibitor liproxstatin-1 (Lip-1) effectively decrease ACSL4, P53 and TRF expression. In vitro, IL-1β induced the reduction of cartilage extracellular matrix expression in mandibular condylar chondrocytes (MCCs). Lip-1 maintain the morphology and function of mitochondria, and inhibited the aggravation of lipid peroxidation and reactive oxygen species (ROS) production which induced by IL-1β. These results suggested that chondrocytes ferroptosis play an important role in the development and progression of TMJOA. Inhibition of condylar chondrocyte ferroptosis could be a promising therapeutic strategy for TMJOA.

**SUMMARY STATEMENT:** Ferroptosis contributed the development and progression of Temporomandibular Joint Osteoarthritis cartilage degeneration. Lip-1 can effective improvement the cartilage degradation of condyle.

## INTRODUCTION

Cell death plays a key role in the development of the body and maintains homeostasis to prevent the development of diseases(Piacentini et al., 2020). A new type of cell death termed, which is a form of programmed cell death driven by iron-dependent lipid peroxidation(Li et al., 2020a). The ferroptosis is characterized by changes in the mitochondrial phenotype, mitochondrial atrophy, and an increase of membrane density(Yu et al., 2017). Ferroptosis results in cytotoxic reactions, damage to mitochondrial function, and cell death(Han et al., 2020). Ferroptosis is proved to be corrected with many degenerative diseases, including Alzheimers, ischemia-reperfusion injury and stroke(Yao et al., 2021). Strikingly, recent study explored that chondrocyte underwent ferroptosis under inflammation and iron overload condition.

Chondrocytes, as the only cell component of cartilage, maintain the integrity of the extracellular matrix (ECM) by balancing the synthesis and degradation of the ECM(Chen et al., 2021b). Chondrocyte injuries is a major event during the progression of osteoarthritis (OA)(Zhang et al., 2021b). Osteoarthritis (OA) is a chronic degenerative disease with progressive features. OA involves structures all parts of the joints in which undergo structural damage and functional imbalances occur as a result of multiple factors(Loeser et al., 2012). D-mannose alleviates anterior cruciate ligament transection (ACLT)-induced OA mouse model by suppressing hypoxia-inducible factor 2α-mediated chondrocyte sensitivity to ferroptosis(Zhou et al., 2021). Iron-overloaded mice exhibit increased cartilage destruction, and intracellular iron uptake is favored in chondrocytes mimicking an OA phenotype(Jing et al., 2020). These findings suggest that ferroptosis is a novel candidate component for development of knee OA progression.

Similar to the knee joint, the temporomandibular joint (TMJ) bears comparable mechanical loading and hence undergoes similar degenerative processes. However, knee articular cartilage and condylar cartilage have different tissue sources, stress modes, and repair and reconstruction methods. TMJOA is a degenerative disease involving articular cartilage, subchondral bone, synovium, ligaments and other tissues around the joints(Liu et al., 2017). The etiology of TMJOA is complex, and the pathological mechanism is not very clear, thus limiting its therapeutic effect. Condylar cartilage is the frontier of the pathogenesis of TMJOA, and the current research focuses on the death of chondrocytes and the degradation of extracellular matrix(Zhang et al., 2022). Programmed cell death such as apoptosis and autophagy play an important role in TMJOA cartilage lesion(Wu et al., 2020). Thus, we proposed that ferroptosis may be related to the occurrence and development of TMJOA cartilage degeneration.

In this study, we found chondrocytes ferroptosis in different TMJOA animal models. Moreover, ferroptosis-specific inhibitor Lip-1 could restore TMJOA cartilage phenotype. It suggested that ferroptosis is closely related to the occurrence and development of TMJOA cartilage lesions.

## RESULTS

### Ferroptosis in Monosodium iodoacetate (MIA) induced- and Interleukin-1 beta (IL-1β) induced TMJOA in vivo and in vitro

To determine the efficacy of MIA and IL-1β on severity and progression of TMJOA, the proteoglycan and structural degeneration of the articular cartilage was evaluated under microscopy by Safranin O-fast green staining and HE staining. It can be observed by HE staining and Safranin O-fast green staining that the lesions in the MIA group and IL-1β are obvious and extensive, mainly manifested as full-thickness structural disorder of cartilage, reduction of chondrocytes in some areas, and thinning of the cartilage layer after 2 weeks (Fig.1A). MIA injection induced serious cartilage matrix destruction and layers disorder. The condylar cartilage lesions were evaluated using the modified Mankin’s score. The results shown that lesions of condylar cartilage were at a relatively early stage after 2 weeks (Fig.1B). MCCs derived from the condylar cartilage (Fig.S1). The MCCs treated with IL-1β (10 ng/ml, 50ng/ml, 100ng/ml) for 24 h. IL-1β decreased the expression of Aggrecan (Acan), type II collagen (COL2), but increase expression of ADAMTS5 and MMP13 in a dose-dependent manner (Fig.1C, D). The effect of 10 ng/ml IL-1β on mandibular condylar chondrocytes (MCCs) has reached cartilage extracellular matrix degradation and elevated inflammation levels. Therefore, subsequent cellular experiments using 10ng/ml IL-1β modeling are credible.

**Fig.1.**
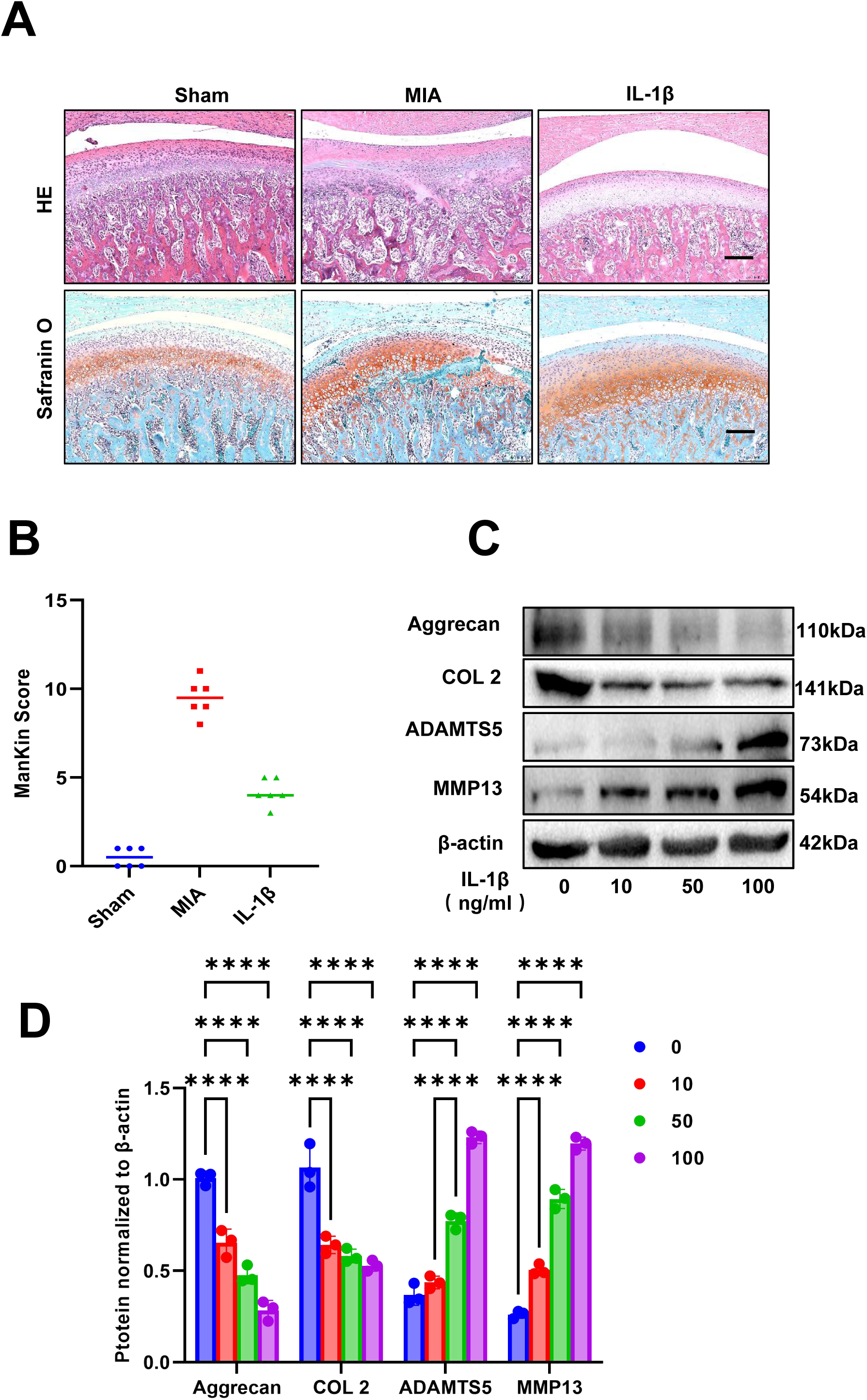
IL-1β and MIA induced chondrocytes injury *in vivo* and *in vitro*. (A) HE staining and Safranin O staining for MIA and IL-1β induced OA models. scale bar = 200 nm. (B) ManKin Score for condylar cartilage OA lesion. N=6. (C)Western blot analysis for protein expression in IL-1β induced chondrocytes injury. (D) Statistical graphs of relative expression protein levels. n=3. ****P<0.0001.

Compared with sham group, the expression of glutathione peroxidase 4 (GPX4) significantly decreased in MIA group and IL-1β group condylar cartilage layer. Meanwhile, Acyl Coenzyme A Synthetase Long Chain Family, Member 4 (ACSL4) and P53 had higher level in MIA and IL-1β groups cartilage layer (Fig.2A). IL-1β treatment decreased the expression of the cellular antioxidant system GPX4. While, IL-1β promoted the expression of lipid peroxidation protein ACSL4, and significantly decreased the expression of transferrin receptor (TFR) protein in chondrocyte (Fig.2C, D). These data revealed that ferroptosis occurred in MIA and IL-1β induced TMJOA animal models.

**Fig.2.**
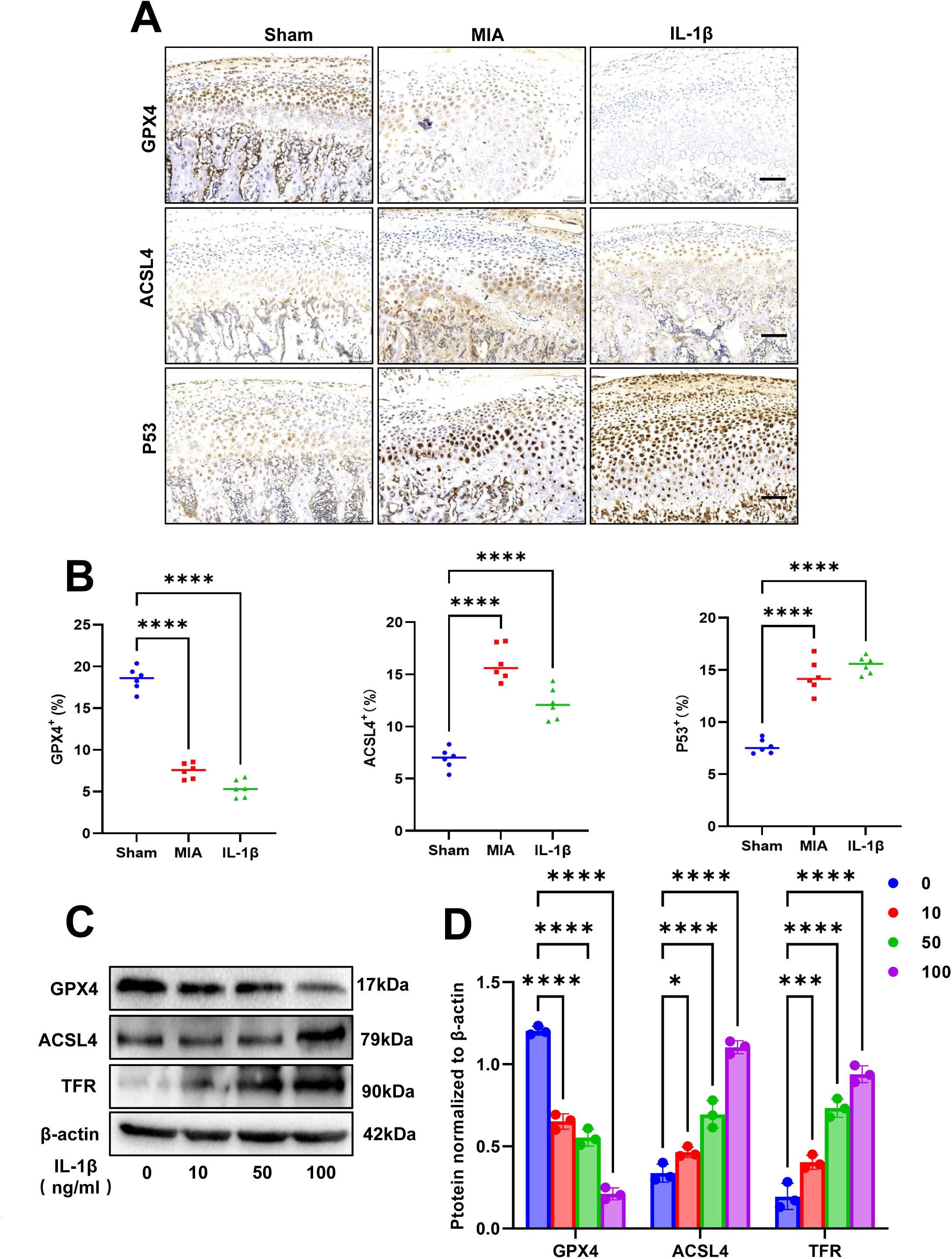
Ferroptosis related protein expressed in IL-1β and MIA induced chondrocytes injury in vivo and in vitro. (A) Immunohistochemical staining of ferroptosis-related proteins expression. N=6. scale bar = 200 nm. (B) Immunohistochemical quantitative results of related proteins in panel A. \*\*\*\**P* < 0.0001. (C) Western blot analysis for ferroptosis protein expression in IL-1β induced chondrocytes injury. (D) Analysis of the relative expression of protein in panel B. \**P*<0.05, \*\*\*\**P*<0.0001. n=3.

### Ferroptosis inhibitor liproxstatin-1 (Lip-1) attenuated ferroptosis related factor expression changes induced by IL-1β

IL-1β induced the morphological and distributional changes of mitochondria. We assessed the prevention of ferroptosis inhibitor Lip-1. The shortened mitochondria or mitochondrial fragments in the IL-1β group were in a scattered distribution. While the mitochondria were prolonged and surrounded the nucleus in the Lip-1 group, those in the Lip-1 lost the normal morphology and the small fragments were heaped in the cytoplasm randomly (Fig. 3A). IL-1β significantly increased the ROS production in the MCCs (Fig.3B). Our results showed that Lip-1 reduced the ROS generation compared with the IL-1β group (Fig.3D). As shown in Fig.2C, an increase in the fluorescence intensity of green JC-1 monomers was observed in the IL-1β group, indicating a significant collapse of mitochondrial membrane potential (MMP, ΔΨm) induced by IL-1β. However, the fluorescence intensity of green JC-1 monomers was weakened in Lip-1 group, which suggested that Lip-1 protected chondrocytes against the loss of ΔΨm induced by IL-1β (Fig. 3E).

**Fig.3.**
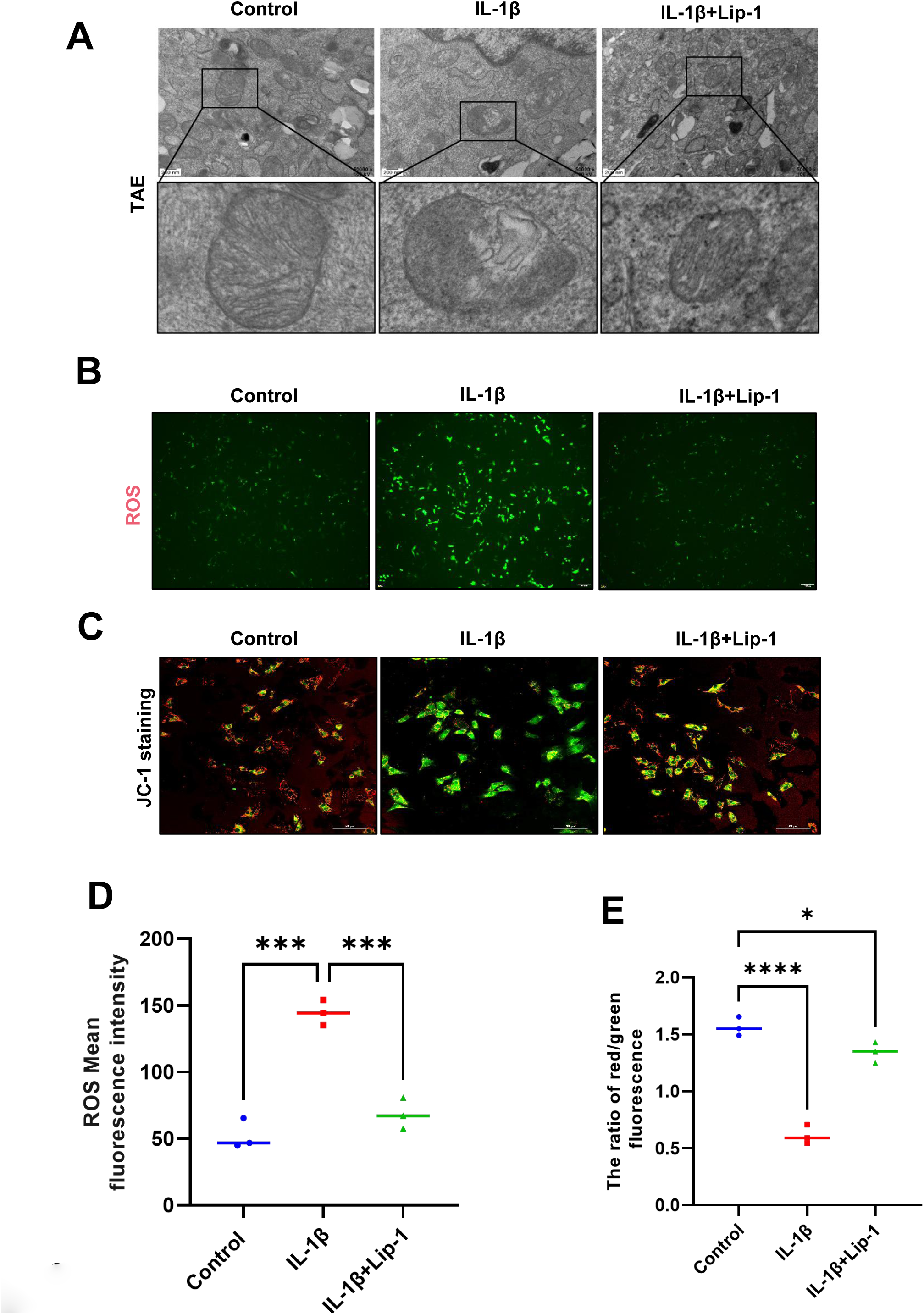
Lip-1 inhibits IL-1β-induced injury in MCCs. (A) Mitochondrial morphology was observed by transmission electron microscopy. scale bar = 200 nm. (B) Detection of intracellular ROS levels using the DAFH-DA probe. (C) Mitochondrial membrane potential was detected using JC-1 probe. (D) Fluorescence intensity quantitative analysis ROS. (E) Fluorescence intensity quantitative analysis ratio results of JC-1 polymer/monomer. \**P*<0.05, \*\*\**P*<0.001, \*\*\*\**P*<0.0001. n=3.

### Lip-1 attenuated reactive oxygen species (ROS), Fe^2+^ and MDA accumulation in MCCs

IL-1β induced Fe^2+^ (Fig.4A) and MDA accumulation in MCCs (Fig.4B) to contribute chondrocyte ferroptosis. Lip-1 significantly reduced level of Fe^2+^ and MDA in chondrocyte (Fig.4A, B). Meanwhile, Lip-1 markedly enhanced the expression of Acan, COL2, but significantly decreased expression of ADAMTS5 and MMP13 in IL-1β induced OA-chondrocytes (Fig. 4C, D). Lip-1 downregulate the expression of ACSL4 in IL-1β induced OA-chondrocytes. The expression of GPX4 increase in Lip-1 treatment OA-chondrocytes. However, we found that Lip-1 had no significant reverse expression of TRF increased by IL-1β (Fig.4E, F).

**Fig.4.**
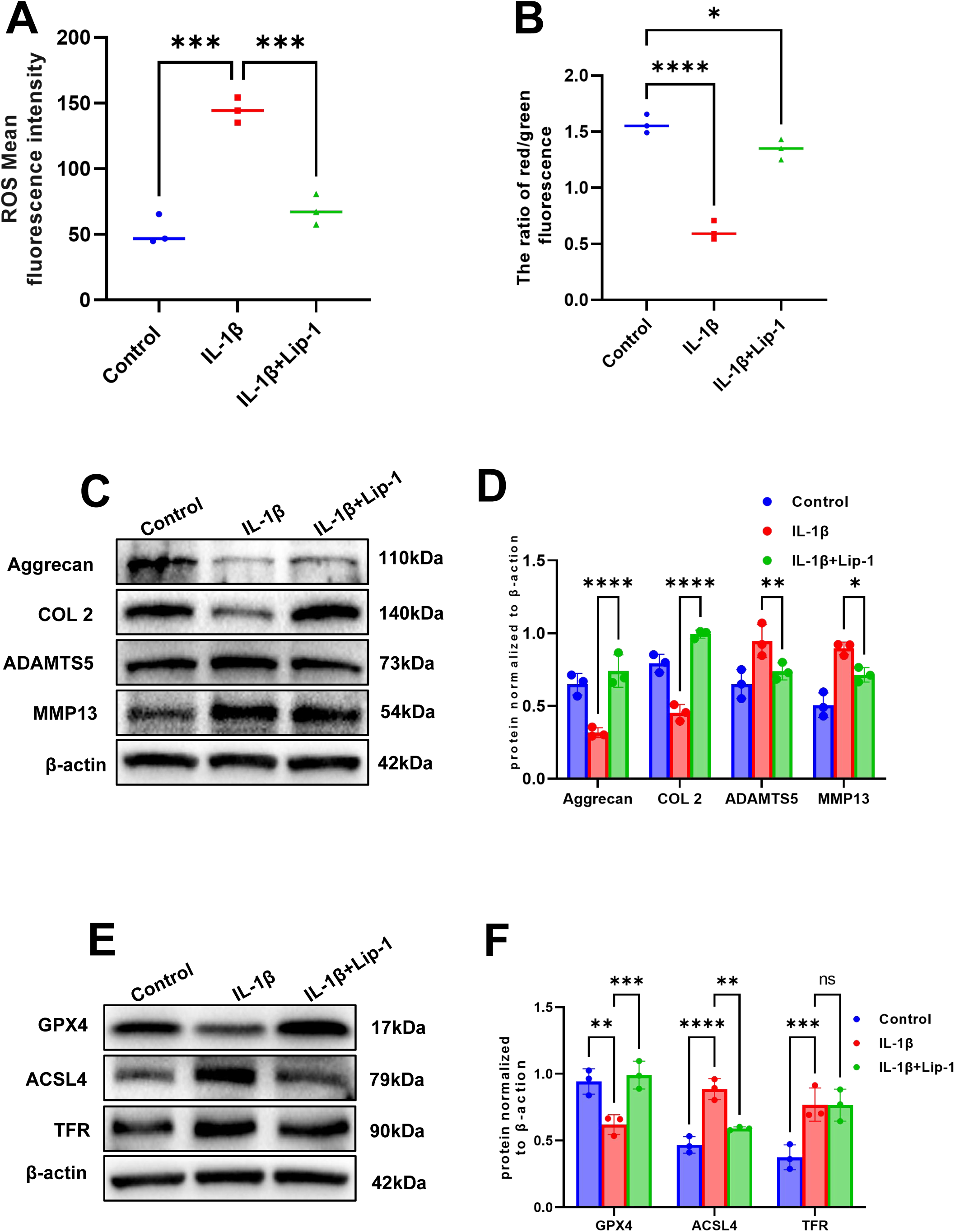
Lip-1 inhibits IL-1β-induced MCCs ferroptosis. (A) Detection of ferrous ion content. (B) Quantitative detection of MDA content. (C) Western blot analysis for protein expression after lip-1 treatment. (D) Analysis of the relative expression of protein in panel C. (E) Western blot analysis for ferroptosis protein expression after lip-1 treatment. (D) Analysis of the relative expression of protein in panel E. n=3.

### Lip-1 improved OA-lesion in IL-1β induced TMJOA models

According to HE staining, Safranin O staining, it was concluded that the Lip-1 injection group had a certain degree of cartilage histological lesions compared with the respective IL-1β induced TMJOA model groups (Fig.5A). The staining results of cartilage extracellular matrix showed that Lip-1 could effectively rescue the decline of the cartilage extracellular matrix Aggrecan and COL2 caused by the TMJOA model (Fig.5B, D). The improved Mankin score was mainly manifested in the normalization of chondrocyte morphology and the increase of extracellular matrix (Fig.5C). The results of immunohistochemical staining found that the expression of ferroptosis-related proteins such as GPX4 was decreased in the respective TMJOA model groups, while the expressions of ACSL4 and P53 were increased, and the Lip-1 group could reverse this change (Fig.5E, F).

**Fig.5.**
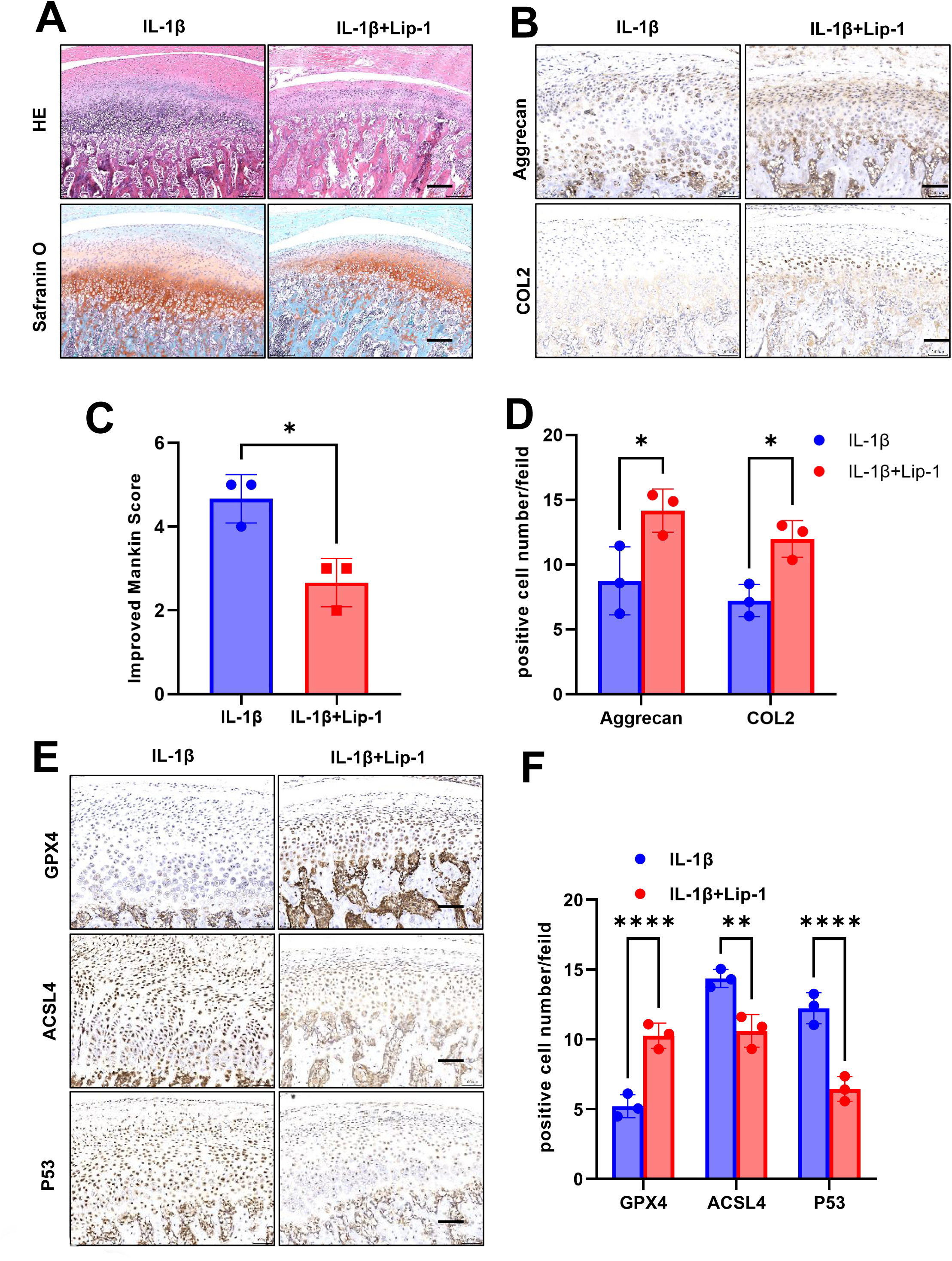
Lip-1 inhibits ferroptosis and protects condylar cartilage in IL-1β-induced OA animal models. N=6. scale bar = 200 nm. (A) HE staining and Safranin O staining of Lip-1 for histological changes. (B) Immunohistochemical staining of cartilage related protein. N=6. scale bar = 200 nm. (C) Modified Mankin’s score of IL-1β induced rat models after Lip-1 treatment. (D) Immunohistochemical quantitative results of related proteins in panel B. (E) Immunohistochemical staining of ferroptosis-related proteins. (F) Quantitative results of related proteins in panel E. *P<0.05, **P<0.01, ***P<0.001, \*\*\*\**P*<0.0001. N=6.

### Lip-1 inhibits ferroptosis and protects condylar cartilage in Occlusion disorder (OD) and unilateral anterior crossbite (UAC) induced TMJOA animal models

There are some differences in chondrocyte responses among different models of TMJOA. In addition to injection, there is extra-articular induction OA. Thus, we used extrarticular induction OA animals to detected chondrocytes ferroptosis. OD induced OA modeling time nodes in this experiment are shown (Fig.S2). After 2 weeks, 4 weeks and 8 weeks modeling, condylar cartilage was analysed histological changes by HE and Safranin O-fast green staining. The chondrocyte layer was obviously thickened, accompanied by the appearance of fissures at the level of the proliferative layer. Safranin O staining showed 4 weeks after OD representing early OA-like manifestations including loss of proteoglycan content, cartilage fibrillation and erosin (Fig.S2B). The score of OD group is near to 4 at 4 weeks, that near to 8 at 8 weeks (Fig.S2C). The expression of GPX4 significantly decreased with the time of OD models establishment in cartilage layer. Meanwhile, ACSL4 and P53 had higher level in OD, compared with control group (Fig.6A). The percentage of positive cells in condylar cartilage was shown (Fig.6B). UAC induced OA modeling time nodes in this experiment are shown (Fig.S3A). The UAC group chondrocyte layer was obviously thickened starting at 2 weeks, accompanied by the appearance of fissures at the level of the proliferative layer, which shown early OA-like manifestations including loss of proteoglycan (Fig.S3B). The Mankin score of UAC group significantly enhanced near to 6 score at 2 weeks, 7 score at 4 weeks (Fig.S3C). The UAC group had obvious cell arrangement disorder, abnormal morphology of the proliferation and hypertrophic layer cells, uneven thickness of the chondrocyte layer and the appearance of fissures in the cartilage layer, and histological lesions were severe.

**Fig.6.**
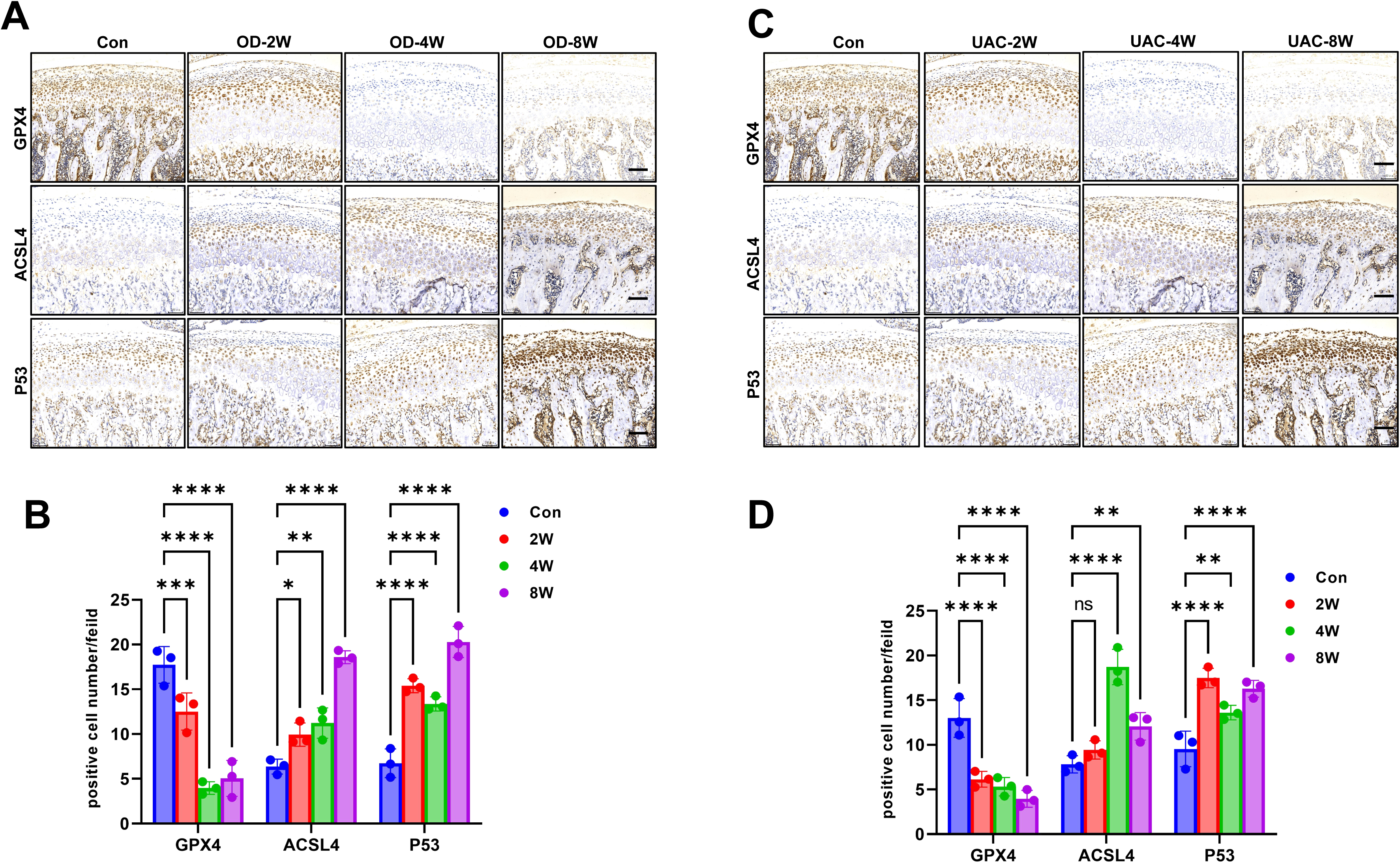
Ferroptosis in condylar cartilage of OD and UAC rat models. (A) Ferroptosis occurs in cartilage layer in the different time periods after OD induced OA animal models. (B) Immunohistochemical quantitative results of related proteins in panel A. (C) Immunohistochemical staining of ferroptosis-related proteins in UAC animal models. scale bar = 200 nm. (D) Immunohistochemical quantitative results of related proteins in panel C. *P<0.05, **P<0.01, ***P<0.001, \*\*\*\**P*<0.0001. N=6. scale bar = 200 nm.

The expression of GPX4 significantly decreased with the time of UAC models establishment in cartilage layer. Meanwhile, ACSL4 and P53 had higher level in UAC (Fig.6C). The percentage of positive cells in condylar cartilage was shown (Fig.6D). After 4 weeks OD modeling, Lip-1 was injected into joint cavity. HE and Safranin O staining shown Lip-1 significantly improved the degradation of condylar cartilage (Fig.7A). Lip-1 significantly improved the degradation of condylar cartilage of UAC models (Fig.7B). Lip-1 decreased the Mankin score of OD animal models (Fig.7C). Meanwhile, Lip-1 decreased the Mankin score of UAC animal models (Fig.7D). Moreover, Lip-1 increased the expression of Aggrecan and COL2 expression in the OD and UAC models cartilage layer (Fig.7 E-G).

**Fig.7.**
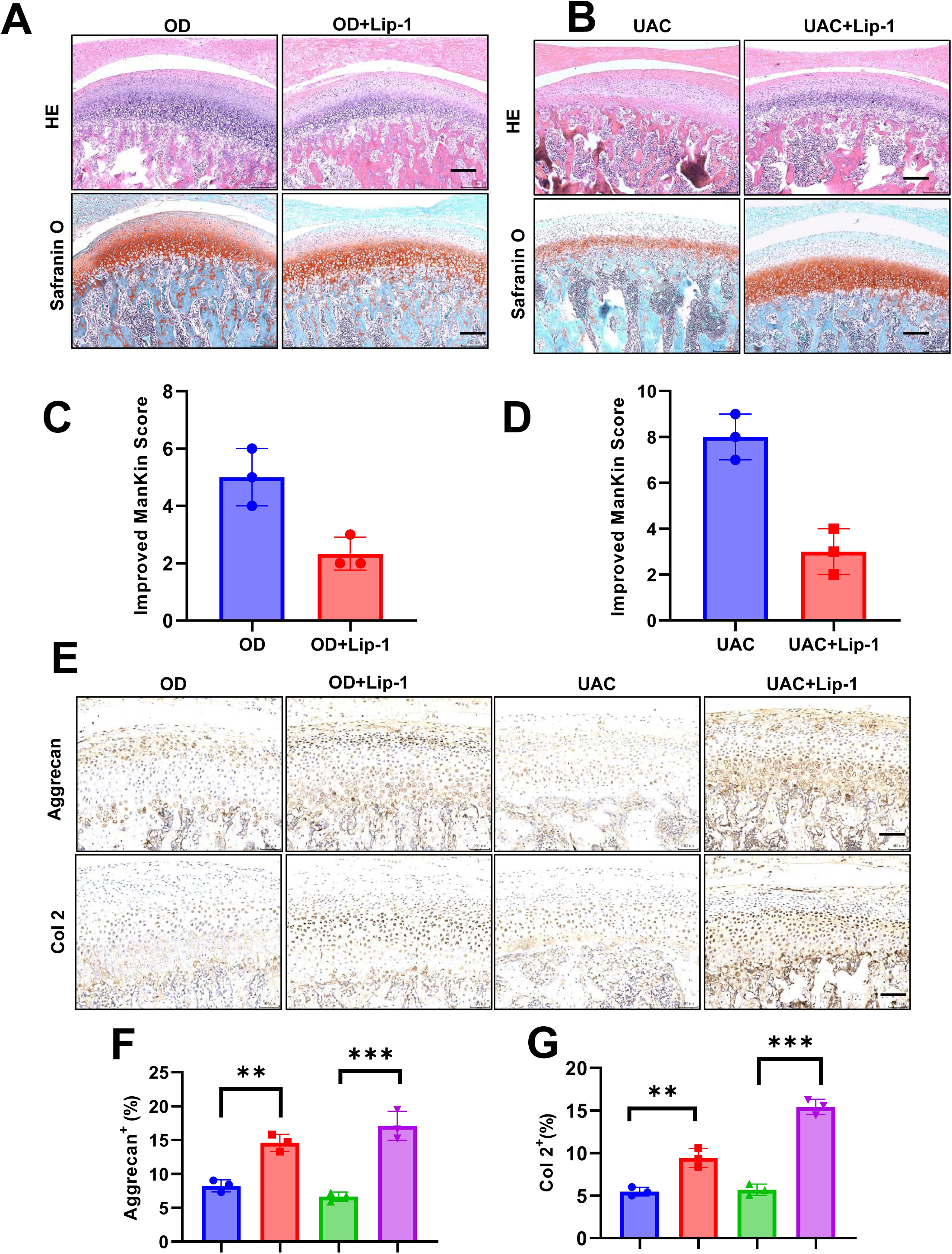
Lip-1 protects condylar cartilage in OD and UAC rat models. (A) HE staining and Safranin O staining of Lip-1 for histological changes OD induced TMJOA rat models. (B) HE staining and Safranin O staining of Lip-1 for histological changes UAC induced TMJOA rat models. (C) Modified Mankin’s score of OD rat models after Lip-1 treatment. (D) Modified Mankin’s score of UAC rat models after Lip-1 treatment. (E) Immunohistochemical staining of cartilage matrix protein. (F) Immunohistochemical quantitative results of aggrecan expression in panel F. (G) Immunohistochemical quantitative results of Col 2 expression in panel F. **P<0.01, ***P<0.005. N=6. scale bar = 200 nm.

In addition to, expression of GPX4 increased, ASCL4 and P53 decreased in OD animal models after Lip-1 injection. Quantitative analysis shown the effects of Lip-1 on level of ferroptosis key protein in condylar cartilage (Fig.8A, B). Lip-1 was injected into joint cavity at 4 weeks in UAC animal models. Moreover, Lip-1 reversed the expression of GPX4, ASCL4 and P53 in cartilage layer in UAC animal models. The level of ferroptosis key protein in condylar cartilage was be quantitatively analyzed (Fig.8C, D).

**Fig.8.**
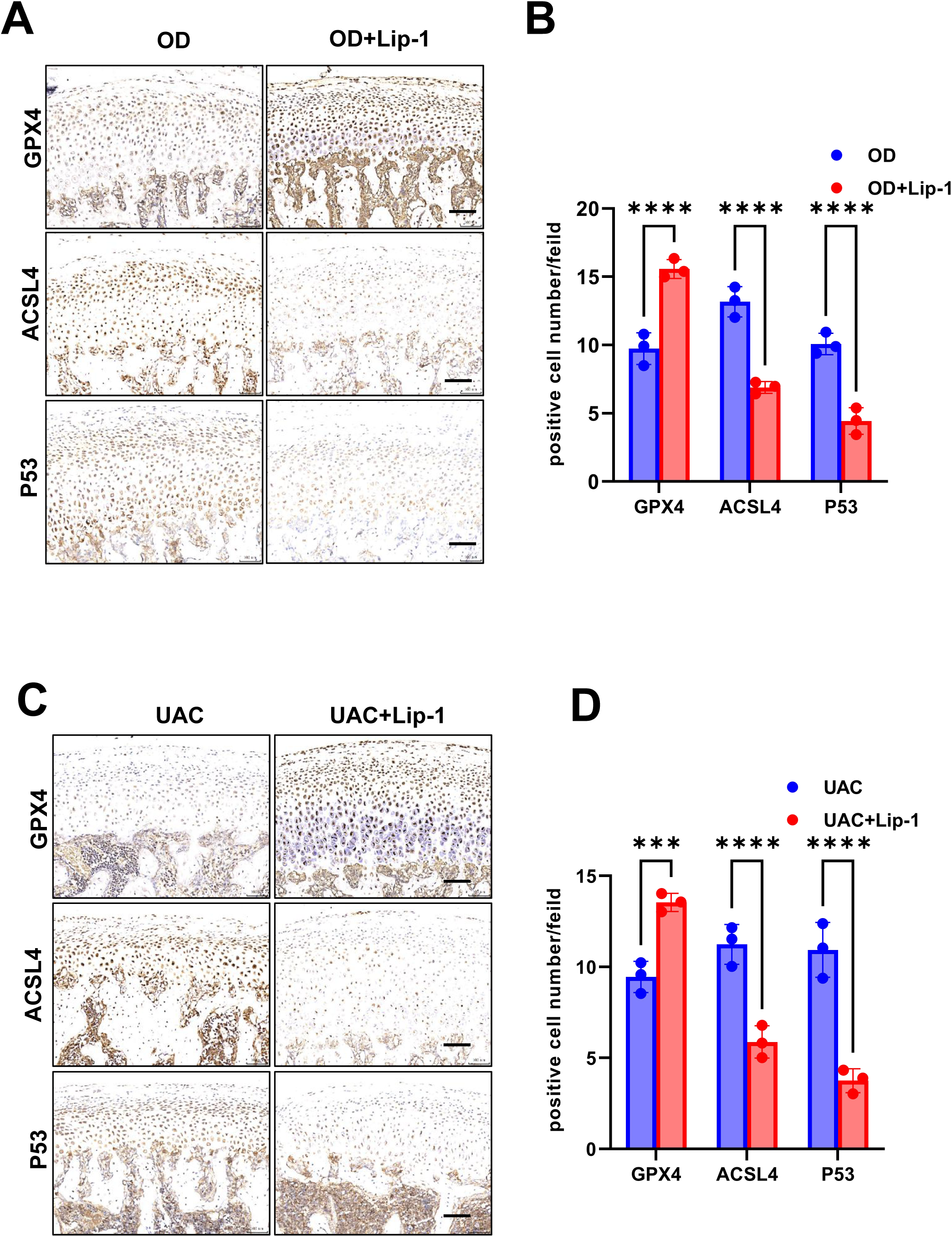
Lip-1 inhibits ferroptosis in OD and UAC rat models. (A) Immunohistochemical staining of ferroptosis protein in OD rat models. (B) Immunohistochemical quantitative results of protein expression in panel A. (C) Immunohistochemical staining of ferroptosis protein in UAC rat models. (D) Immunohistochemical quantitative results of in panel C. ***P<0.005, \*\*\*\**P*<0.0001. N=6. scale bar = 200 nm. SFig.1

## DISCUSSION

Chondrocyte injuries is a major event during the progression of OA(Rim et al., 2020). It’s well studied that chondrocyte injuries can be attributed to the cell death including necrosis, apoptosis, ferroptosis and autophagy(Musumeci et al., 2015, Kapoor et al., 2011, Zhou et al., 2022). However, it is still unknown whether condylar chondrocytes undergo ferroptosis contribute to TMJOA lesion.

Animal model will lay the foundation of the serial study of TMJOA(Chen et al., 2022). TMJOA animal models can promote the exploration of the pathogenesis, development, treatment, and prevention of this disease (Na et al., 2021). Therefore, we established TMJOA models of four different lesions by injecting monosodium iodoacetate (MIA) or IL-1β into the articular cavity, and unilateral anterior crossbite and occlusion disorder inducing. MIA-induced animal model of OA is well-known for reproducing the symptoms as in the case of OA patients (Li et al., 2021, Liao et al., 2020). When injected into the joint cavity, MIA disrupts the glycolytic energy metabolism in chondrocyte and causes cell death, leading to inflammation and cartilage damage, which cause severe OA phenotype in a short period of time (Feng et al., 2019). IL1β plays a crucial role in the pathogenesis of OA. Thus, IL-1β was used as one of the common methods for OA modeling in vitro (Liu et al., 2022). Occlusion disorder is a key factor which contribute to TMJOA in clinic(Kalladka et al., 2022). OD models mimic the pathogenesis of human TMJOA to the greatest extent by raising the occlusion of the unilateral maxillary molars of SD rats to achieve continuous occlusal interference (Zheng et al., 2018). Moreover, the UAC model achieves biomechanical stimulation of condylar cartilage by interfering with the occlusion of the rat anterior teeth(Zhou et al., 2020). UAC induced TMJOA animal models was most widely studied. Our data demonstrated that the four models induced degradation and destruction of condylar cartilage to different degrees. Among the four animal models in this experiment, the MIA model has the most severe cartilage lesions, reaching the middle and late stages of osteoarthritis. The observation results of the comparison between the OD model and the UAC model in the same period showed that the OD model was in the early stage of the development of cartilage lesions (Liu et al., 2018), while the UAC model was in the middle stage of the development of the cartilage according to Mankin score and histological staining (Li et al., 2020b).

In our four different TMJOA animal models, we found ferroptosis related protein expressed in cartilage layer. Ferroptosis is a form of lipid peroxidation-induced cell death that can be regulated in a number of ways (Yang and Stockwell, 2016). The level of lipid peroxidation was detected the dynamic balance of the antioxidant and peroxidative systems of tissue cells (Maiorino et al., 2018). GPX4, as important antioxidant enzymes in cells, inhibit lipid peroxidation and resist the occurrence of ferroptosis P53 can inhibit the transcriptional activity of SLC7A11, resulting in the inhibition of antioxidant system function (Li et al., 2008). P53 can promote the expression of SAT1, and its downstream targets can be involved in promoting the production of lipid reactive oxygen species (Sun et al., 2022). In addition, ACSL4 is also a key enzyme in promoting lipid peroxidation(He et al., 2022). Both P53 and ACSL4 can promote the production of lipid peroxidation and accelerate the occurrence of ferroptosis(Chen et al., 2021a). We observed decreased expression of GPX4 and increased expression of P53 and ACSL4 in the cartilage layer of the TMJOA rat models. These results confirmed that antioxidant function inhibition and the enhancement of peroxidative reaction related to OA lesion development.

Lip-1 was mainly manifested to inhibit lipid peroxidation(Feng et al., 2019). Normal articular cartilage has no blood supply, it is difficult to deliver the effective drug concentration of Lip-1 to the articular cartilage through blood circulation if the drug is administered by conventional intraperitoneal injection(Huang et al., 2022). Therefore, in this study, Lip-1 was injected into the temporomandibular joint cavity, and the dose was slightly adjusted compared with the conventional Lip-1. Intraperitoneal injection reduced the concentration and prevented adverse reactions due to high local drug concentration in the cartilage. However, no Lip-1 concentration gradient was set for comparative observation in this experiment, so follow-up experiments are still needed to ensure the minimum effective drug concentration for Lip-1 injection treatment.

The Lip-1 could significantly reduce the cellular divalent iron ion level again (Zhang et al., 2021a). Lip-1 could indirectly cause a decrease in divalent iron ion levels (Feng et al., 2019). In vivo experiments, it was observed by histological staining that Lip-1 improved cartilage destruction in different TMJOA models by increasing the content of cartilage extracellular matrix, increasing the number of chondrocytes and normalizing the cell arrangement. However, the effect of Lip-1 on the reversal of cartilage lesions varied among the models. In the OD model, where the lesion was relatively early, Lip-1 treated the chondrocyte number and morphology relatively best, followed by the IL-1β and UAC models. While in the middle and late stages of OA destruction, such as the MIA model, where the cartilage lesion was more severe, Lip-1 could only improve this condition to a certain extent. This suggests that early intervention of TMJOA cartilage lesions has the best effect, while intervention of mid- to late-stage TMJOA can only salvage the cartilage pathology to a certain extent, and there are limitations to fully restore it to normal levels.

Overall, our study demonstrates chondrocyte ferroptosis related to OA lesion development. The inhibitor of ferroptosis Lip-1 significantly improved the cartilage degradation in OA animal models and chondrocyte injures induced by IL-1β. It suggested that ferroptosis is a potential mechanism for TMJOA.Lip-1 may be used as an active regent for treatment of TMJOA.

## MATERIAL AND METHODS

### TMJOA Rat models

8 wk-old male Sprague-Dawley (SD) rats (weighing160 to 180g) were randomly divided into normal (Con, n=6), sham-operated control (Sham, n=6) and experimental groups (n=10 rats/group). In experimental groups, monosodium iodoacetate (MIA, 40mg/kg, I2512, Sigma-Aldrich) and IL-1β (10000 ng/ml, TP750014, ORIGENE) injected the right cavity of TMJ respectively, sham group was injected the equivalent volume of PBS. Occlusion disorder (OD) was created by abnormal dental occlusion force based on previously reported(Zhang et al., 2022). The occlusal of the rat left maxillary first molar was raised by tying the occlusal ring with a corrective ligation wire (0.2mm diameter) and placing it on the occlusal surface. The unilateral anterior crossbite (UAC) was used to establish unilateral anterior crossbite prosthesis to induce TMJOA. Ferroptosis inhibitor liproxstatin-1 (Lip-1, 1mg/kg, S7699, Selleck) was injected into bilateral TMJ joint after each model was established.

All animal procedures were approved by the Ethical Review Committee of Animal Experiments, Kunming Medical University and it is prior to the initiation of the study (Approval number: kmmu20211222). This study was performed in strict accordance with this approved protocol and adhered to the ARRIVE guidelines (Animal Research: Reporting In Vivo Experiments) for conducting animal research.

### Histological Staining and Scoring

The mandibular joint of rats in each group was removed as a whole, fixed with 4% paraformaldehyde, decalcified with 15%EDTA, dehydrated with gradient alcohol, continuous sagittal section with a thickness of 4μm, and hematoxylin eosin staining (HE staining) and Safranin O staining (G1371, Solarbio) were performed to evaluate the degree of tissue lesions. TMJOA scores were evaluated by three oral pathologists using an improvement Mankin’s score, which was expressed as mean ± standard deviation (SD).

### Immunohistochemical Staining

Immunohistochemical (IHC) staining was performed using a standard protocol. Sections were incubated overnight at 4℃ with the following primary antibodies: Glutathione Peroxidase 4 (GPX4, ab125066, 1:200, Abcam), Acyl Coenzyme A Synthetase Long Chain Family, Member 4 (ACSL4, ab155282, 1:200, Abcam), P53 (ab33889, 1:200, Abcam), Aggrecan (Acan, ab36861,1:200, Abcam), type II collagen (COL 2, ab34712, 1:400, Abcam). All sections were incubated with a biotinylated secondary antibody and stained using the R&D HRP-DAB staining kit (R&D Systems, USA) and then counterstained with hematoxylin. The positive cells were analysed by Image J software (NIH, USA).

### Statistical Analysis

Statistical analysis was accomplished using Graphpad Prism 8 software (GraphPad company, San Diego, CA, USA). Data are expressed as the means ± standard deviation for each group. Comparisons between groups were evaluated by unpaired two-tailed Student’s t test between two groups or by one-way ANOVA followed by Tukey’s test for multiple comparisons. P<0.05 were considered as significant difference between groups.

### Isolation and chondrocyte culturing

Mandibular condylar chondrocytes (MCCs) were isolated from the condylar cartilage of 3-day-old Sprague-Dawley rats. Briefly, condylar cartilage tissue was minced and digested with 0.25% trypsin (Gibco) for30 min and 0.25% collagenase II (Invitrogen) at 37°C for 6 h sequentially. Chondrocytes were collected and cultured in an incubator containing 5% CO_2_ at 37°C with Dulbecco’s DMEM/F12 (HyClone) supplemented with 10% fetal bovine serum (FBS, Gibco) and 100 U/ml penicillin G sodium, 100 μg/ml streptomycin sulfate (Gibco). The MCCs were collected and passed to P3 expansion for subsequent experiments. The culture medium was replaced every 2 days during the incubation period. The MCCs were treated with IL-1β (10ng/ml, 50ng/ml, 100ng/ml, TP750014, ORIGENE) for 24 hours.

### Western Blot analysis

Protein extraction and Western blot analysis were performed as the protocol. Membranes were probed with anti-GPX4(ab125066, 1:1000, Abcam), ACSL4 (ab155282, 1:1000, Abcam), P53 (ab33889, 1:1000, Abcam), Aggrecan (Acan, ab36861,1:1000, Abcam), type II collagen (COL 2, ab34712, 1:1000, Abcam), Transferrin Receptor (TFR, ab269513, 1:1000, Abcam), ADANTS5 (ab41037, 1:1000, Abcam), MMP13(ab39012, 1:1000, Abcam). β-actin (1:1000, 20536-1-AP, Proteintech), followed by incubation with secondary horseradish peroxidase-conjugated antibody (Santa Cruz Biotechnology, Santa Cruz, CA, 1/5000). After incubation with primary and secondary antibodies, immunoreactive bands were visualized using β-actin served as an internal standard for semi-quantification. Protein blots were visualized using Western ECL Substrate Kit (Thermo Pierce) and exposed using a ChemiDoc XRS+ system (BD, Franklin Lakes, NJ). The intensity of bands was quantified by Image J software

### Transmission electron microscope (TEM)

TEM is an important method for the characterization of mitochondrion as it can directly visualize their inner architecture. In brief, the cells were washed with PBS and resuspended, and then post-fixed by 2% OsO4 (Electron Microscopy Sciences), dehydrated with different concentrations of alcohol and acetone, and then embedded in epoxyresin. Ultrathin sections were stained with uranyl acetate (E. Merck) and lead citrate (Sigma-Aldrich). A TEM (JEM-100CXII) was used for observation.

### Mitochondrial membrane potential (MMP, ΔΨm) measurement

Mitochondrial membrane potential was measured by the fluorescent probe 5,5′,6,6′-tetrachloro-1,1′,3,3′-tetraethylbenzimidazolocarbocyanine iodide (JC-1) detection kit (68-0851-38, Thermo) according to the manufacturer’s instructions. JC-1 accumulates in mitochondria matrix with high membrane potential, resulting in the formation of JC-1 aggregates and the emission of red fluorescence. When the ΔΨm is low, the JC-1 monomer is formed and yields green fluorescence. In brief, the cells were stained with JC-1 staining solution in the dark at 37°C. After incubation for 30 min, the cells were washed twice with JC-1 staining buffer and immediately observed with a fluorescence microscope. To quantify the MMP, cells were prepared as mentioned above and resuspended with JC-1 staining buffer, and finally analyzed using a microscope (Olympus, Japan).

### Evaluation of intracellular reactive oxygen species

Levels of ROS were measured using fluorescent probe reactive oxygen species assay kit (Thermo, C6827) as per the manufacturer’s instructions. Briefly, the cells were stained with 10 μM DCFH-DA (C6827, Invitrogen) in the dark at 37°C. After 30 min of incubation, the cells were washed twice with serum-free DMEM media and immediately visualized with a fluorescence microscope (Evos flauto; Life Technologies).

### TBA fluorescence

Lipid peroxidation was detected by the MDA detection kit (TBA fluorescence method, Solarbio, BC0025) according to manufacturer’s instructions. Briefly, cells supernatant was transfered 100 μL of sample into microcentrifuge tube in duplicate. Prepare 50 μM MDA standard by add 50 μL MDA standard into 950 μL dH2O, then following the table to generate 50, 30, 15, 5 and 0 μM MDA standards. Add 100 μL 10% TCA into each sample and standard tubes, vortex and incubate for 5 minutes at room temperature. This acid treatment clarifies the samples by precipitating interfering proteins and other substances, and also catalyzes the TBARS reaction. Pre-heat water both or heat block to 95-100 °C. Carefully remove 100 μL (in duplicate) reaction mixture into a 96-well clear microplate. Read the optical density at 535 nm.

### Iron Assay

The Iron Assay Buffer in this kit has an acidic pH that enables this release of iron/ferric ions into the solution. The Iron Assay Kit (Abcam, ab83366) was used for measuring ferrous (Fe2^+^) and/or ferric (Fe3^+^) iron in biological samples. According the protocol output immediately on a colorimetric microplate reader at 593 nm.

## Acknowledgments

We would like to thank Hefeng Yang for providing funding support.

## Competing interests

No competing interests declared.

## Funding

This research was funded by the “Ten Thousand Talents Program” Young Top Talents of Yunnan Province, Yun Ren She Tong [2019] No. 206 from Hefeng Yang. Natural Science Foundation of Yunnan Province (202001AY070001-083) from Yu Hu.

## Data availability

Raw data has been uploaded to Figshare (DOI: 10.6084/m9.figshare.22329793).

## CRediT Authorship Contribution Statement

Bei Cheng: Conceptualization, Formal analysis, Investigation, Writing – original draft preparation, visualization

Jun Zhang: Conceptualization, Formal analysis, Investigation, Writing – original draft preparation, visualization

Qinhao Shen: Formal analysis, Investigation, Writing – review and editing

Zheyi Sun: Software, Formal analysis, Investigation, visualization, Writing – review and editing

Yinwei Luo: Writing – review and editing

Yu Hu: Conceptualization, Resources, Writing – review editing, Funding acquisition, Supervision, Project administration

---

**Fig.S1.** MCCs culturing and indentification of characteristics. (A-B) is cellular morphology. (C) MCCs toluidine blue staining. (D) Col II staining of MCCs.

**Fig.S2.** OD induced TMJOA animal models. (A) OD modeling visual diagram. (B) HE and Safranin O staining for OD animal models cartilage. (C) Mankin Score for OD animal models.

**Fig.S3.** UAC induced TMJOA animal models. (A) UAC modeling visual diagram. (B) HE and Safranin O staining for UAC animal models cartilage. (C) Mankin Score for UAC animal models.

## Notes

### Competing Interest Statement

The authors have declared no competing interest.

